# Numerical simulation for behind armor blunt trauma of human torso under non-penetrating ballistic impact

**DOI:** 10.1101/586594

**Authors:** Fan Tang, Zerong Guo, Mengqi Yuan, Xinming Qian, Zhiming Du

## Abstract

A human torso finite element model with high bio-fidelity was developed to study the behind armor blunt trauma (BABT) of pistol cartridge on human torso with bulletproof composite structure (BCS) and the effect of buffer layer (expandable polyethylene, EPE) on BABT. The bulletproof structure was made of multilayered composite of aluminum alloy (AlSi_10_Mg) and thermoplastic polyurethanes (TPU), and the ANSYS/LS-DYNA software was used to simulate the blunt ballistic impact process of pistol cartridge on human torso. Results indicated that the BCS could resist the shooting speed of 515 m/s without being broken. During the process of pistol cartridge shooting the BCS, the energy of pistol cartridge was transmitted to the human organs through the BCS, thereby causing human injury. Moreover, the mechanical response parameters of various organs were determined by the distance between the human organs and the impact point. The sternal fracture and liver rupture were not produced based on the threshold stress of sternum and liver injury, no matter whether the buffer layer was added or not. According to the Axelsson injury model, a slight to moderate injury was created when there was no buffer layer, but the injury level was trace to slight caused by the buffer layer with thickness of 1.0 mm, and the buffer layer with thickness of 2.5 mm and 5.0 mm caused subtle BABT. It was concluded that the buffer layer could effectively reduce the BABT, and the reduction was related to the thickness of the buffer layer. This study reveals the mechanism of the BABT, which can provide a theoretical basis for the design of the bulletproof structure and the evaluation of structural bulletproof performance and protection performance.

## 1. Introduction

The body armor is a kind of equipment used to protect the bullets or fragments from harming the human body, which plays an extremely important protective role for reducing the casualties of soldiers and police [1-3]. Although the body armor can effectively reduce the penetrating injury caused by the bullets or fragments, the energy of pistol cartridge will be transmitted to the human through the body armor, causing injury to the thoracic and abdominal organs, even as indirect brain injury, this non-penetrating injury phenomenon is called BABT [4, 5]. Among American soldiers, there have been cases of death due to BABT. Although the body armor remains intact after attack, the broken ribs stabbed into heart leading to death [6].

Body armors are categorized by bulletproof materials into soft body armor, rigid body armor and soft-rigid composite body armor [7]. The soft body armor has the advantages of softness, lightweight and comfort, but it is greatly deformed under the shooting of pistol cartridge, so the BABT caused by soft body armor is serious [8, 9]. The rigid body armor has excellent anti-penetration performance and non-penetration injury performance, but its disadvantages are rigid, bulky and uncomfortable [10, 11]. Compared with the bulletproof structure of a single material, the soft-rigid BCS combines the advantages of soft bulletproof structure and rigid bulletproof structure, and has better anti-penetration performance and non-penetration injury performance [12-14]. According to GA 141-2010 “Police ballistic resistance of body armor” [15], the backing materials (such as clay) are used to simulate the human torso to assess the bulletproof performance of body armor. In the case of a valid hit for bullet, the body armor can block the warhead, and the maximum backface signature is less than or equal to 25 mm, that is, the body armor can effectively protect the killing effect of the pistol cartridge, while the NIJ (National Institute of Justice) standard is 44 mm [16]. It is impossible to ascertain the injury situation of the human wearing the body armor, so the evaluation method is only suitable for evaluating the bulletproof performance of body armor, and it is also difficult to be applied to the evaluation of the protective performance of body armor to the human.

Many scholars have also done a lot of research on BABT, but mainly focused on the research of animal experiments and soft bulletproof materials such as fibers, and the human model used in previous numerical simulation is not realistic. In 1969, Shepard [17] reported the first case of BABT of lung, which started the study of BABT. Roberts [18-20] has established a human torso finite element model with high fidelity, and assessed the BABT caused by NIJ (National Institute of Justice) soft body armor by testing the mechanical response parameters of the organ near the ballistic impact point. Kunz et al. [21] studied the heart injury caused by BABT, and analyzed the formation and propagation of pressure waves in the heart. Sonden et al. [22] studied the blunt trauma of landrace pigs protected by a ceramic/aramid body armor, and proposed that trauma attenuating backing can reduce the BABT. Zhang et al. [23] found that high velocity BABT generates high pressure and acceleration in the animal’s spine, and pressure wave is an important factor in causing BABT. The numerical simulation results of Wickwire et al. [24] showed that both sternum displacement and pressure are sensitive to impact energy and location of projectile.

The BCS made of AlSi_10_Mg and TPU was taken as the research object, and a real human torso model was developed. The ANSYS/LS-DYNA finite element analysis software was used to simulate the mechanical response of human organs under the blunt ballistic impact of pistol cartridge.

## 2. Finite element model

### 2.1. Human torso model

Human torso model consisted of skin (epidermis, dermis and hypodermis), skeleton (sternum, rib, costal cartilage, vertebral column, intervertebral disc, clavicle and scapula), internal organs (kidney, pancreas, liver, heart, lung, spleen and stomach), blood, diaphragm and muscle, the half section view of human torso model as shown in Fig. 1(a). The skeleton, internal organs, blood, diaphragm and muscle model were modeled as tetrahedral elements and the Lagrange approach was used. The epidermis, dermis and hypodermis were divided into Belytschko-Tsay shell elements with thicknesses of 0.1 mm, 2.0 mm and 2.0 mm, respectively. The contact between human organs was set to automatic single surface contact other than the skin, and the contact between skin and muscle was defined as tied contact. The material models and material parameters of human organs [18, 19, 25] were given in Table 1.

**Fig. 1.**
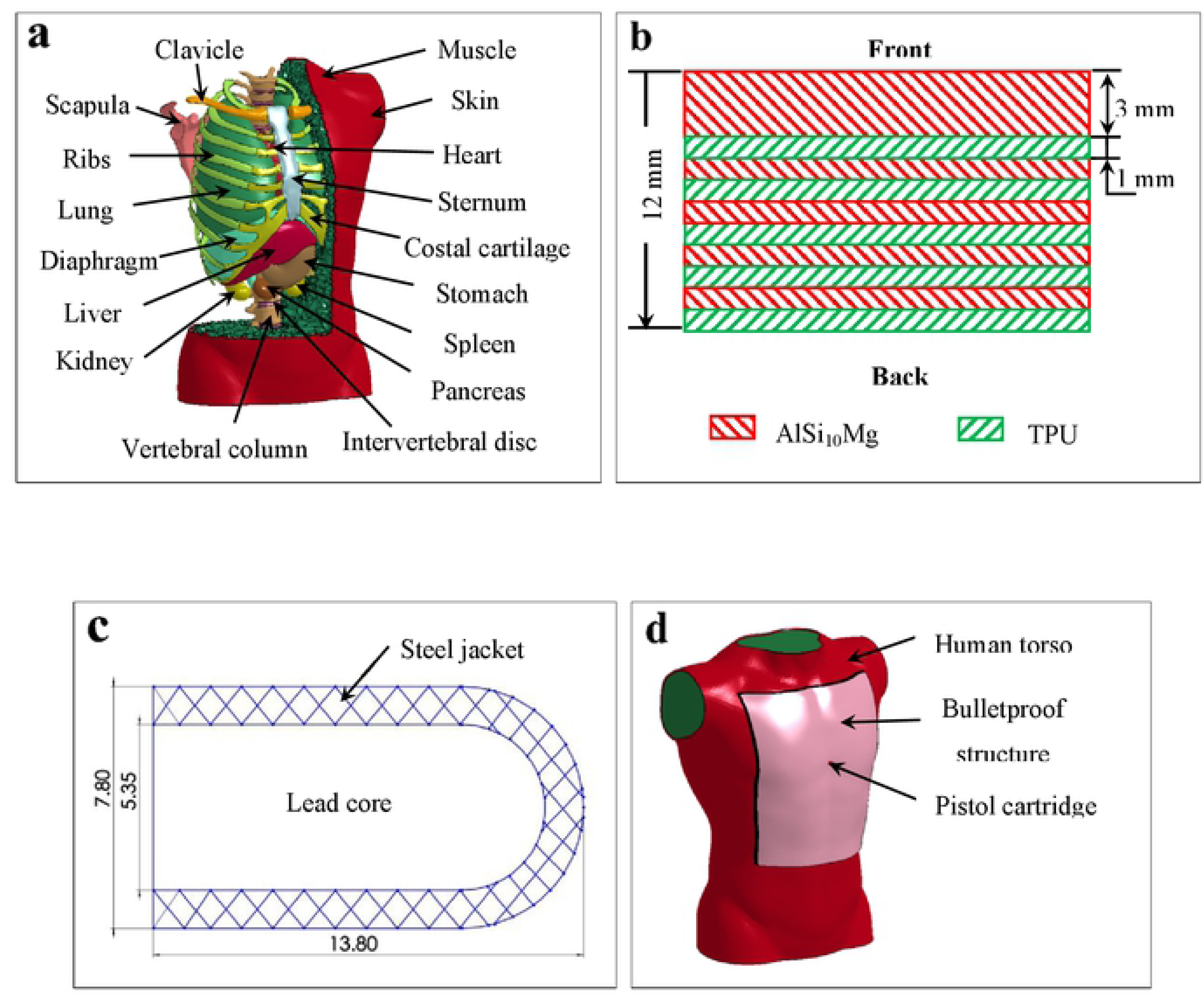
Finite element model including: (a) human torso, (b) bulletproof composite structure, (c) pistol cartridge structure and (d) calculation model.

**Fig. 2.**
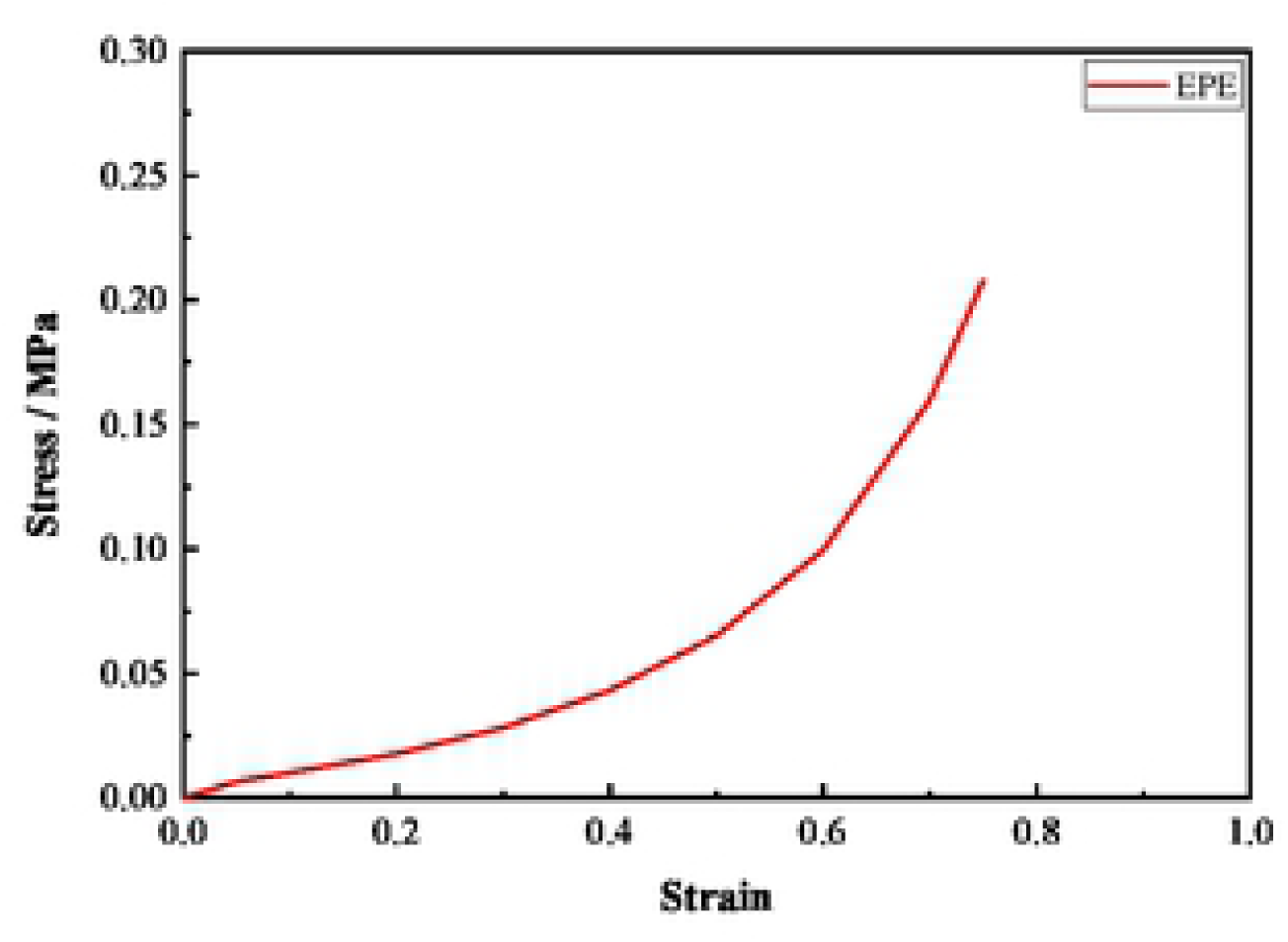
Stress-strain curve of EPE.

### 2.2. Pistol cartridge and BCS model

The BCS was made of “rigid-soft-rigid” composite of AlSi_10_Mg and TPU with a total thickness of 12 mm. The composite method of bulletproof structure was shown in Fig. 1(b). Both AlSi_10_Mg and TPU were divided into 5 layers, except that the thickness of the front face was 3 mm, the thickness of the remaining layers was 1 mm, the protective area was 0.11 m^2^, and the areal density was 2.37 g/cm^2^. Type 51 pistol cartridge was used to shoot BCS, the mass of warhead was 5.6 g, the initial velocity was 515±10 m/s, the structure of warhead was round, and the material was steel jacket and lead core, which geometry was shown in Fig.1(c). The expandable polyethylene (EPE) buffer layer was added between human torso and BCS to weaken the blunt impact of BCS on human torso, and the effect of the thickness of the buffer layer on BABT was studied. The material models and material parameters of pistol cartridge [26], AlSi_10_Mg [27], TPU [28] and EPE [29] were given in Table 2∼5, respectively.

The finite element model (FEM) was built in g-cm-μs unit. The pistol cartridge, BCS and buffer layer were divided into hexahedral elements, as well as Lagrange approach was adopted. The contact between steel jacket and lead core, pistol cartridge and BCS, pistol cartridge and buffer layer, pistol cartridge and human torso were defined as erosion surface-to-surface contact. Moreover, the contact between AlSi_10_Mg layer and human torso, TPU layer and human torso, buffer layer and human torso were set to automatic single surface contact, and the contact between AlSi_10_Mg layer, TPU layer and buffer layer were defined as tied contact. To prevent the influence of stress waves generated by the impact of pistol cartridge on the calculation results, non-reflective boundary conditions were imposed at the boundary of BCS and buffer layer [30]. The calculation model was shown in Fig. 1(d).

### 2.3. Validation of FEM

In the Roberts’s research [18], the 9 mm Full Metal Jacket (FMJ) Luger ammo with an initial velocity of 430 m/s was used to shoot Kevlar soft body armor with the thickness for National Institute of Justice (NIJ) Level IIIa vests of 10.82 mm. The diameter of the warhead was 9.03 mm, the weight was 8 g, the structure was oval, and the material was lead core and full metal jacket. The structure of 9 mm FMJ Luger ammo was shown in Fig. 3. The pistol cartridge and body armor models used by Roberts and the human torso model established in this paper were used for the validation of FEM, the material parameters of pistol cartridge and body armor were shown in the reference [10, 31].

**Fig. 3.**
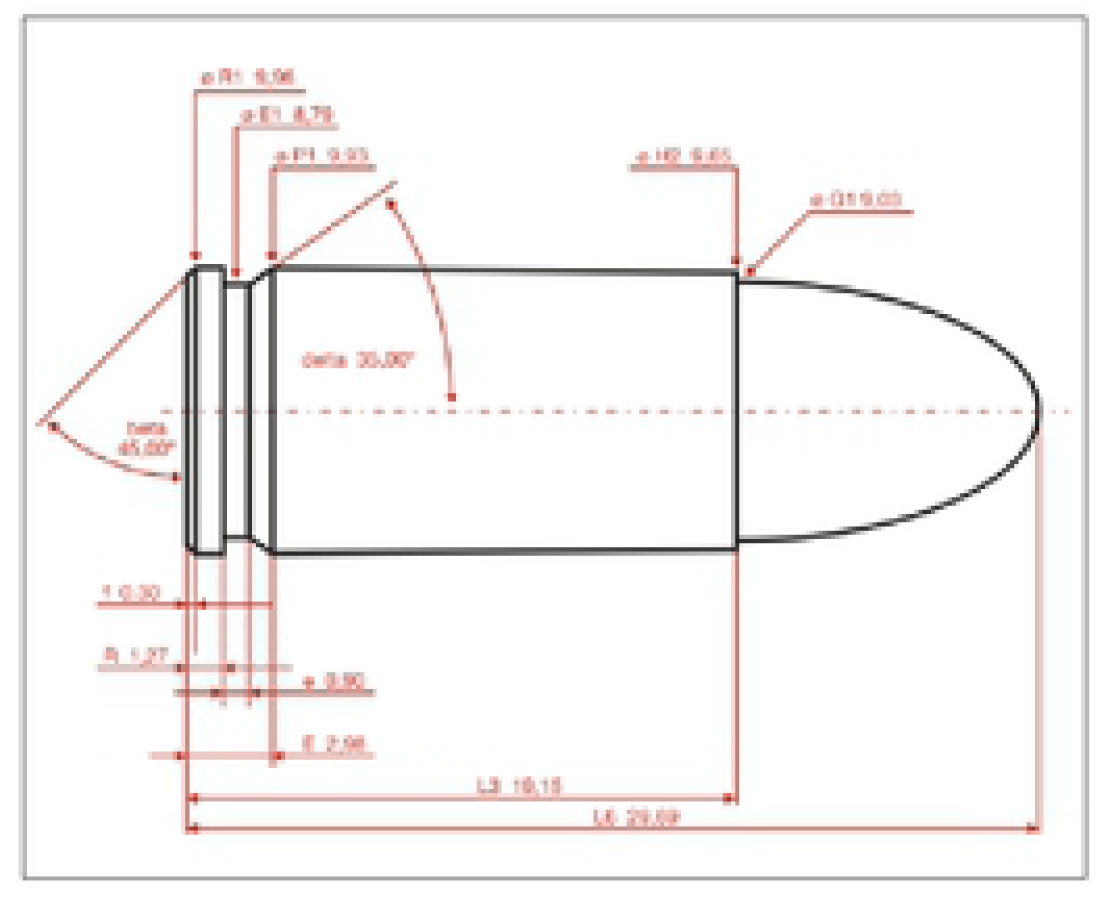
The structure of 9 mmFMJ Luger ammo.

Fig. 4 shows the simulated and Roberts’s pressure curve at the middle of the front of heart. The peak pressure of heart obtained by numerical simulation is 0.958 MPa, while the peak pressure of heart given by Roberts is 0.743 MPa, which the relative error is 28.9%. It is concluded that the numerical simulation result is in good agreement with Roberts’ research, so the finite element model established in this paper is reasonable. However, the trend of the pressure curve still differs, which is caused by the difference between the human torso model, body armor and measurement points.

**Fig. 4.**
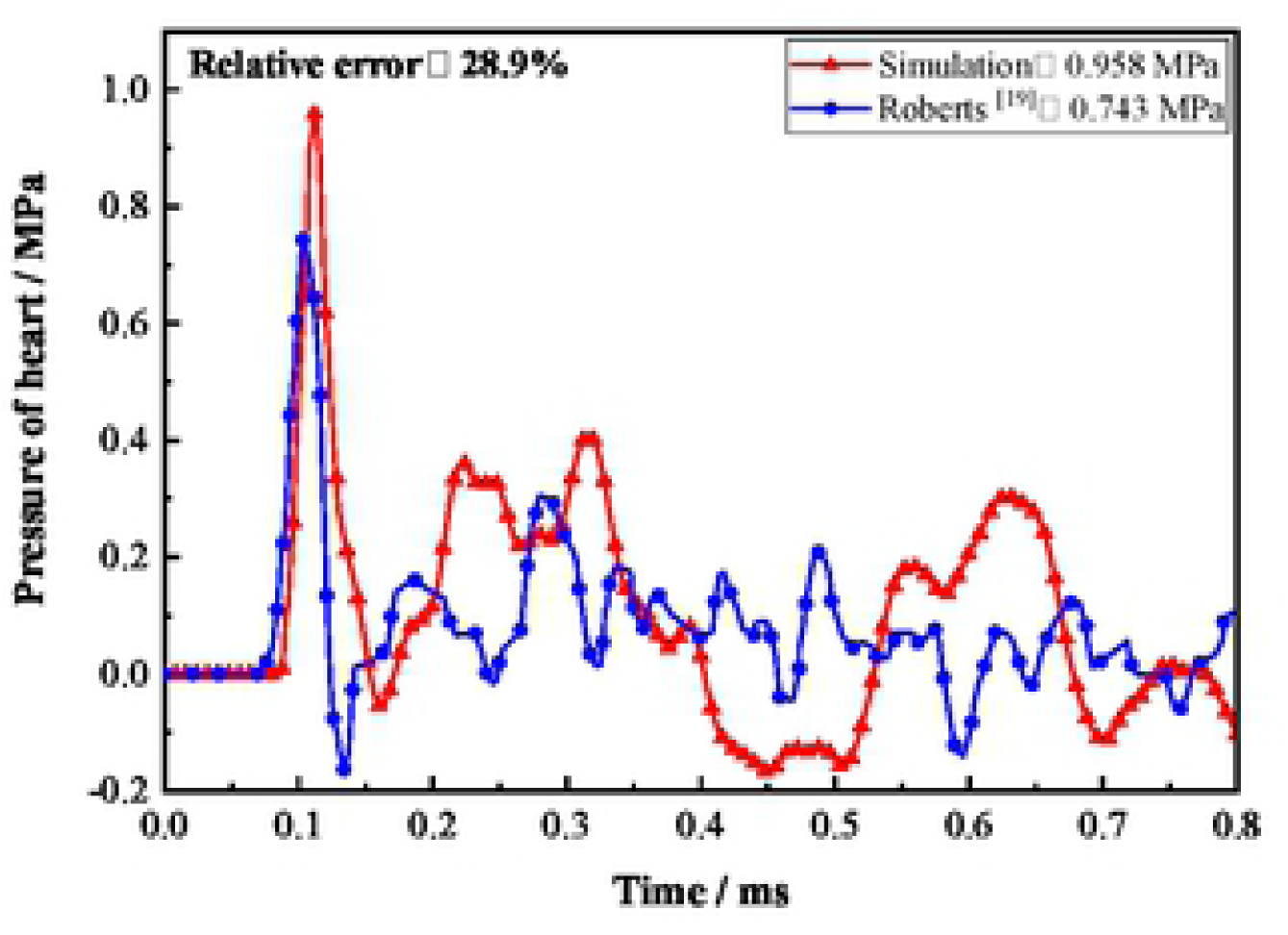
Comparison of heart pressure obtained by simulation and Roberts.

## 3. Results and discussion

The pistol cartridge shot the human torso at an initial velocity of 515 m/s (initial kinetic energy was 742.63 J) from the middle of sternum in front of heart. The bulletproof performance of BCS, the mechanical response of human organs, and the effect of buffer layer on BABT were obtained by numerical simulation.

### 3.1. Bulletproof performance of BCS

Fig. 5 shows the impact process of pistol cartridge on BCS, Fig. 6 reveals the changes of the velocity of pistol cartridge, total energy of BCS and human torso. The positive velocity of pistol cartridge drops to zero at 68 μs, followed by inverse motion, with a maximum inverse velocity of 60.74 m/s. The reason for the inverse motion of pistol cartridge is that TPU is an elastomer rubber, as well as skin and muscle have some elasticity. After the positive velocity of pistol cartridge decays to zero, the energy of BCS and human torso will be transferred to the pistol cartridge due to the rebound of pistol cartridge, TPU, skin and muscle, so the pistol cartridge begins to move in inverse after gaining kinetic energy. The maximum total energy absorbed by BCS is 208.00 J, accounting for 28% of the initial kinetic energy of pistol cartridge, while the maximum total energy delivered to the human torso is 37.02 J, which accounts for 5% of the initial kinetic energy of pistol cartridge. It is obvious that the BCS absorbs more energy than human torso. On one hand, the BCS is directly shot by pistol cartridge, so it bears most of the energy of pistol cartridge. On the other hand, the pistol cartridge does not directly shoot the human torso, and the last layer of BCS is TPU, which acts as a buffer, thereby reducing the blunt ballistic impact of BCS on human torso. Consequently, the BCS absorbs more energy and transmits less energy to human torso.

**Fig. 5.**
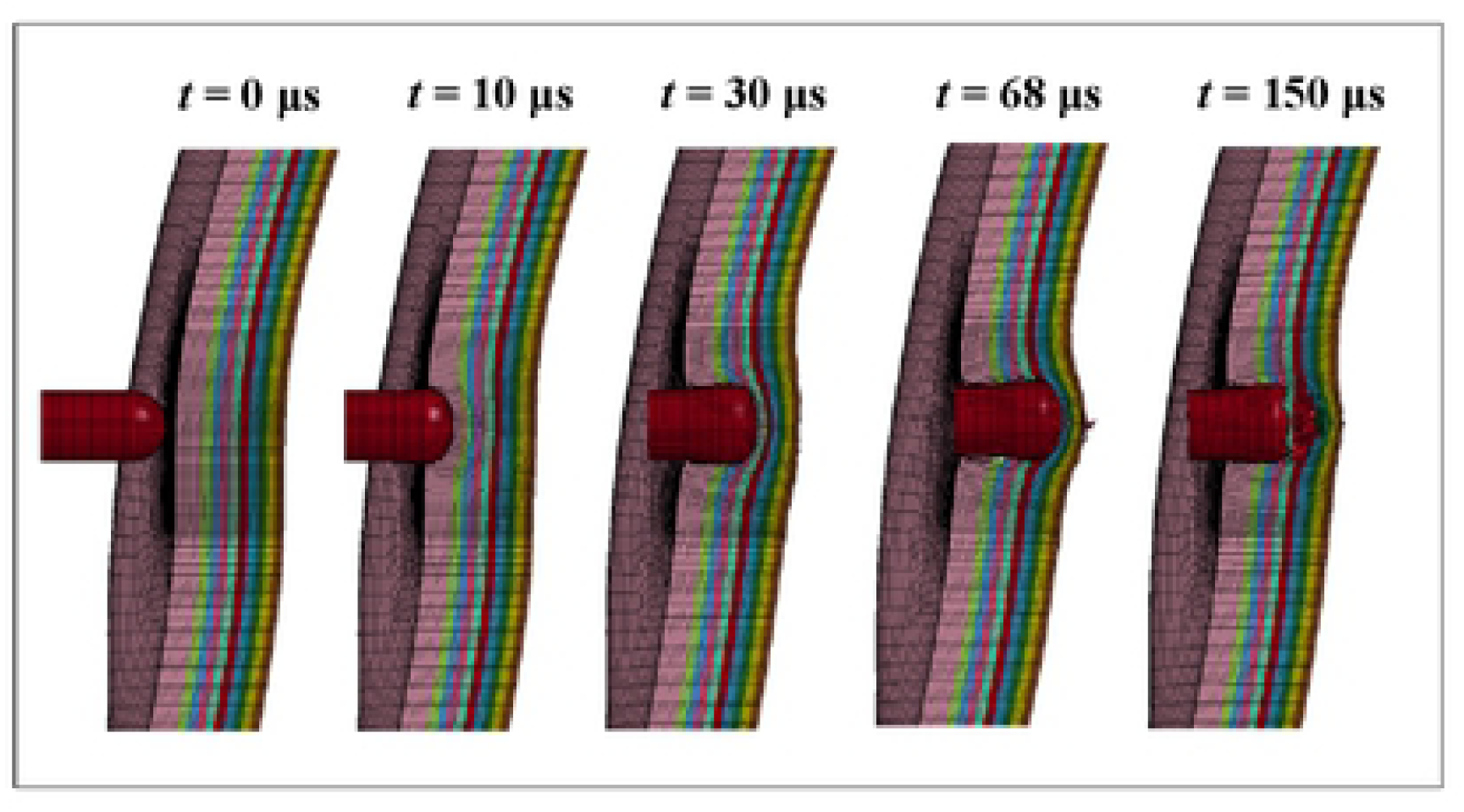
Impact process of pistol cartridge on bulletproof composite structure.

**Fig. 6.**
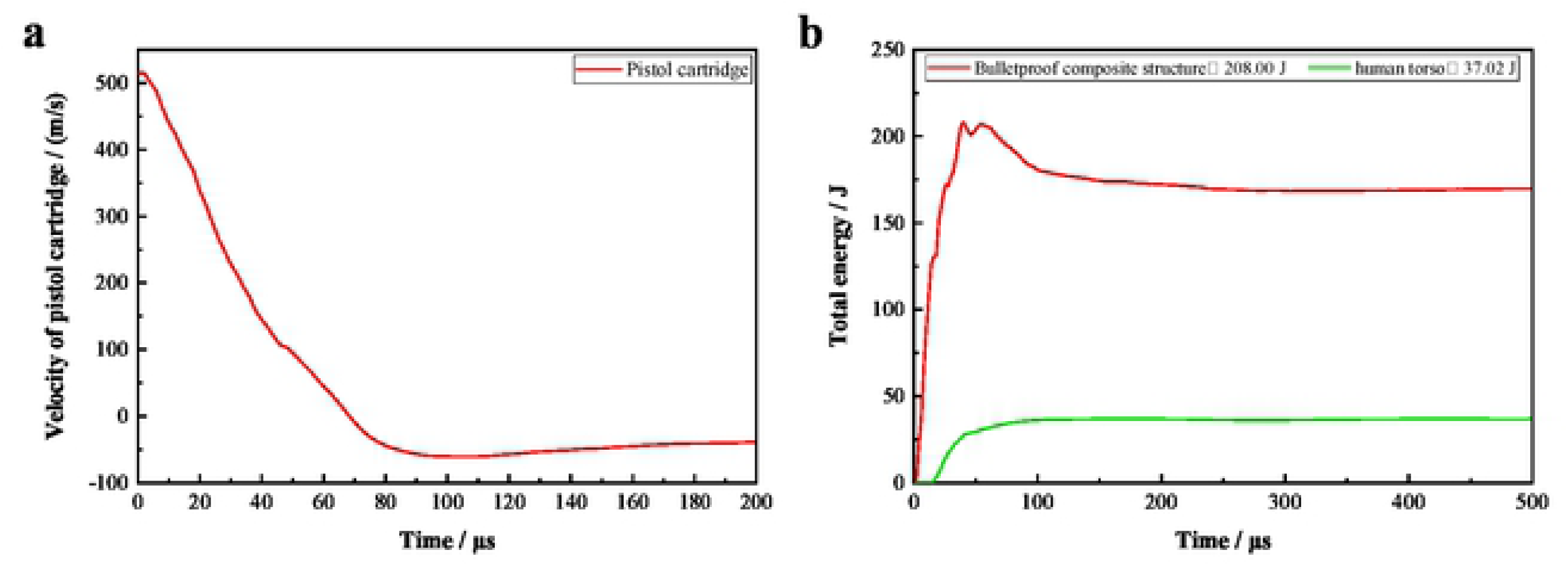
Time histories of (a) velocity of pistol cartridge and (b) total energy of bulletproof composite structure and human torso.

It is observed from Fig. 5 that when the BCS is subjected to the impact of pistol cartridge, it will be deformed, sunken or even broken. The deformation of the AlSi_10_Mg layer is mainly fragmentation, while the TPU layer mainly suffered from compression deformation. The protective mechanism of BCS mainly relies on the deformation or fragmentation of BCS to absorb part of the kinetic energy of pistol cartridge and converts it into its own internal energy and kinetic energy, and transmits part of the energy of pistol cartridge to human torso at the same time.

The numerical simulation results indicate that the BCS is not broken down by pistol cartridge, and the maximum depression depth of skin is 4.98 mm, which is less than 25 mm specified by the Chinese standard [32] and 44 mm specified by the NIJ standard [16]. Hence, the pistol cartridge does not directly shoot the human torso. Meanwhile, it can be found that the depression depth of skin behind the BCS is small, the reason is that the human torso has a certain supporting effect on BCS, especially for skeleton, its stiffness and damping are relatively large [33].

### 3.2. Blunt ballistic impact process of pistol cartridge on human torso

The BCS is not penetrated by pistol cartridge, so the pistol cartridge dose not directly hit the human torso. However, when the pistol cartridge interacts with BCS, 5% of the initial kinetic energy (37.02 J) of pistol cartridge is transmitted to the human organs through BCS and spreads around the impact point, thereby causing human organs injury.

Fig. 7∼11 displays the distribution of stress and pressure of skin, heart, lung, liver and skeleton under the blunt ballistic impact of pistol cartridge. The stress first appears in front of skin, skeleton, lungs, heart and liver, and the time of appearance is 12 μs, 14 μs, 20 μs, 34 μs, 40 μs, respectively. It is found that the mechanical response time of various organs has certain differences under the influence of the distance between the human organs and impact point. The stress wave and pressure wave of lungs are generated at 20 μs, and gradually spread to the whole lung at 200 μs, 1000 μs, 2000 μs, which reflects the propagation of stress waves and pressure waves from surface to inside and from near to far. Under the blunt ballistic impact of pistol cartridge, the skin at the impact point is the first to appear stress and the peak stress is the largest, then the stress spreads around the impact point and gradually spreads through the muscles to the skeleton and internal organs, such as sternum, lungs, heart and liver. As the stress wave propagates in the human organs, its energy will be attenuated, resulting in the most serious injury to the human organs near the impact point, while the human organs injury far from the impact point is the lightest. For example, the maximum stress in front of lungs is 12.79 kPa, and the maximum stress in the back of lungs is 1.95 kPa.

**Fig. 7.**
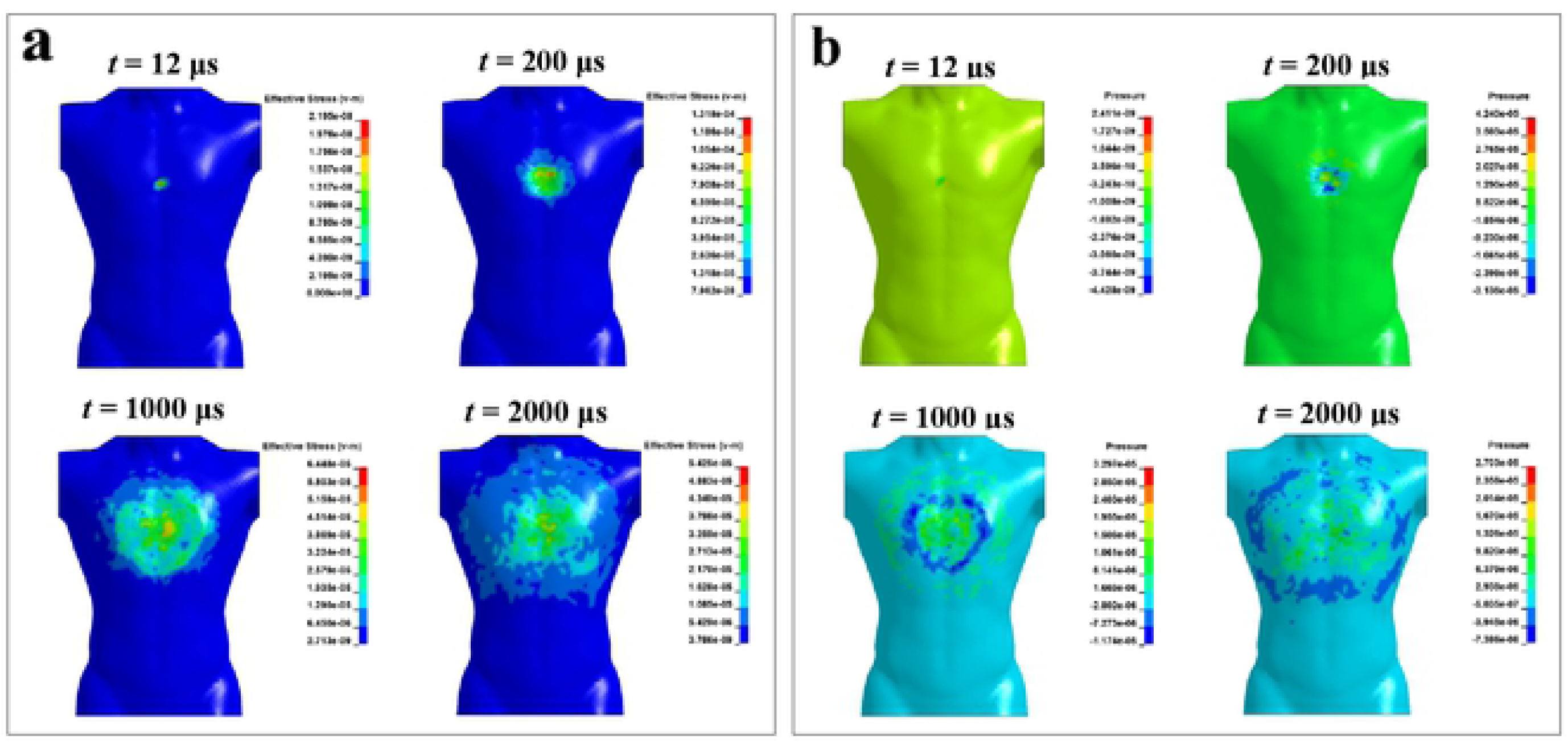
Distributions of (a) stress and (b) pressure of skin at different time.

**Fig. 8.**
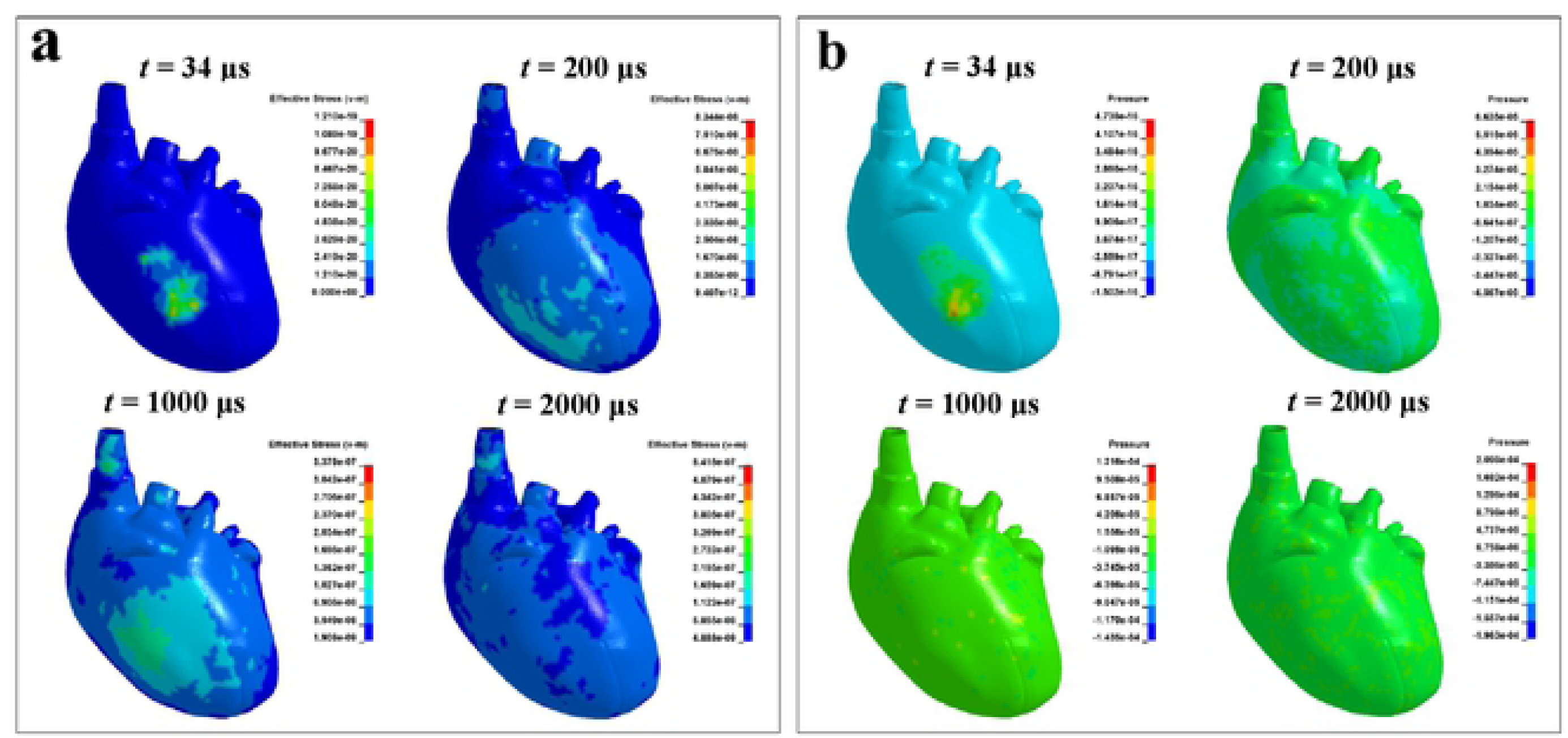
Distributions of (a) stress and (b) pressure of heart at different time.

For the same organ, the propagation laws of stress waves and pressure waves have extremely significant differences. At 20 μs, 200 μs, 1000 μs, and 2000 μs, lung stress is mainly concentrated in front of lungs, and pressure has spread throughout the lungs at these times, as shown in Fig. 9. At 200 μs, the lung stress is positive, and the maximum stress and minimum stress are different by six orders of magnitude, while the pressure has both positive and negative values, which differ by two orders of magnitude. During the blunt ballistic impact of pistol cartridge on human torso, part of the energy of pistol cartridge is transmitted to the human organs in the form of pressure waves. A high-pressure shock wave is formed in front of pistol cartridge, its speed is close to the speed of sound in the human organs, and the peak can reach tens of atmospheric pressure [34, 35]. Consequently, the pressure wave propagates faster and has a wider range of influence. In contrast, the stress can only occur when the human organs is deformed by external force, so its propagation speed is slow and the incidence is small [36].

**Fig. 9.**
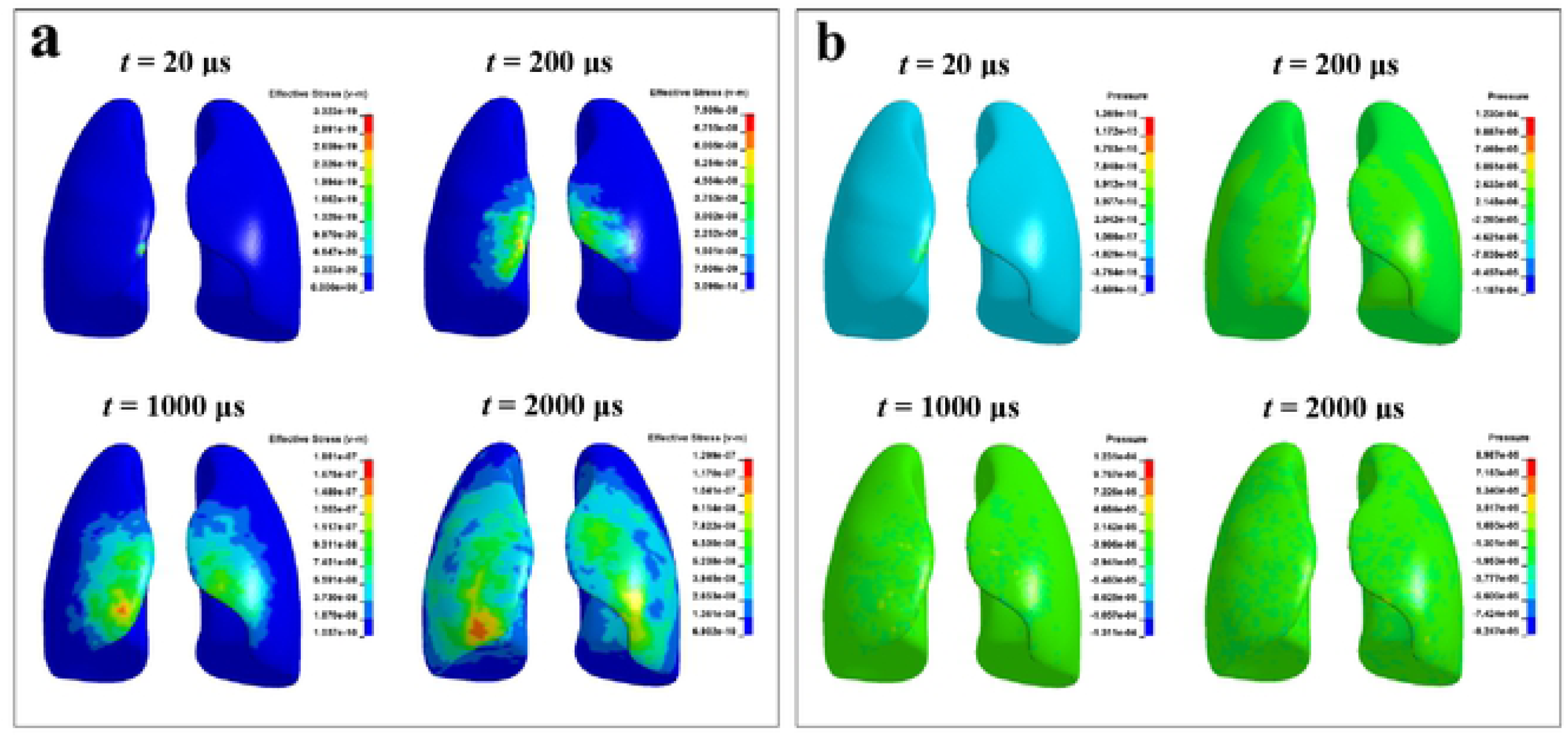
Distributions of (a) stress and (b) pressure of lung at different time.

**Fig. 10.**
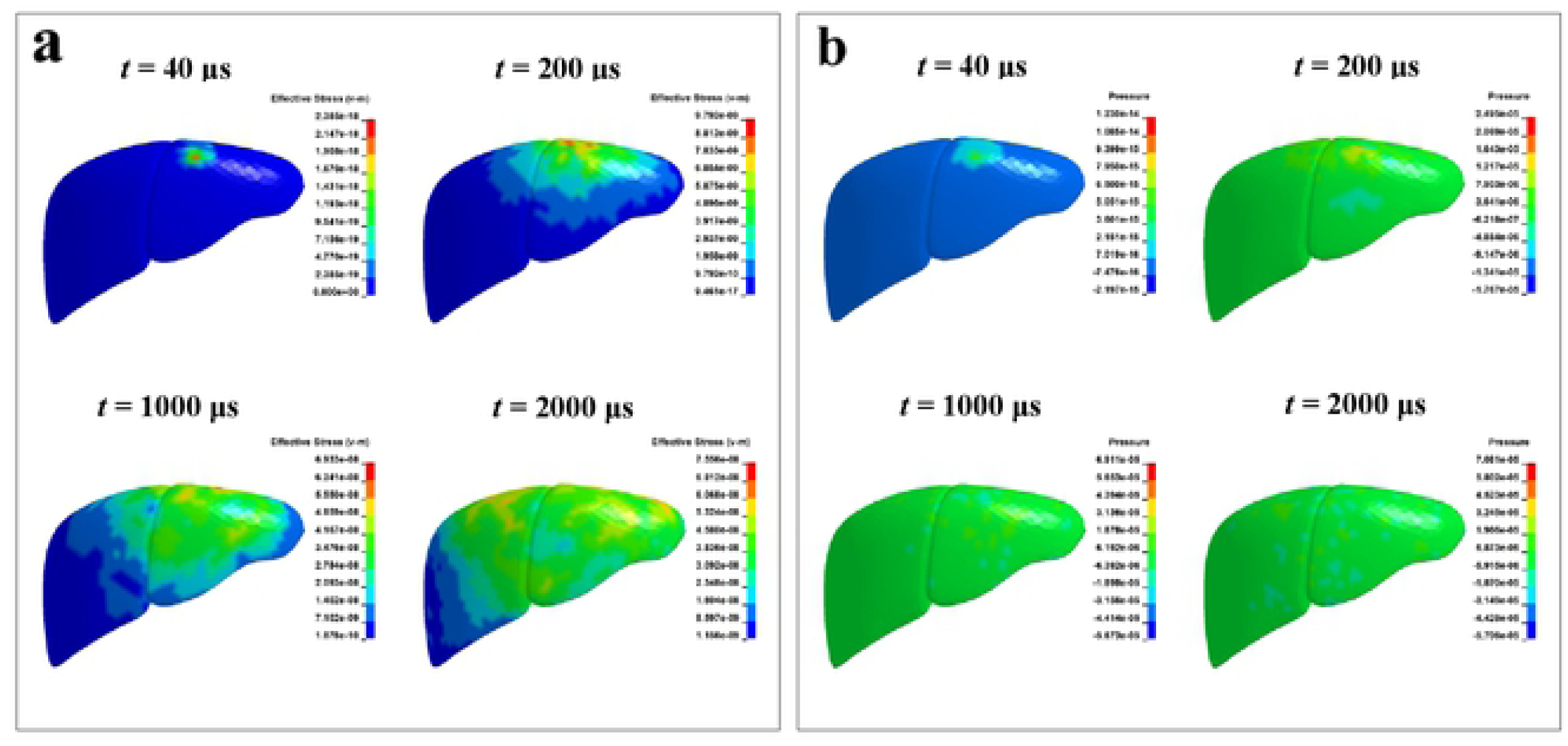
Distributions of (a) stress and (b) pressure of liver at different time.

**Fig. 11.**
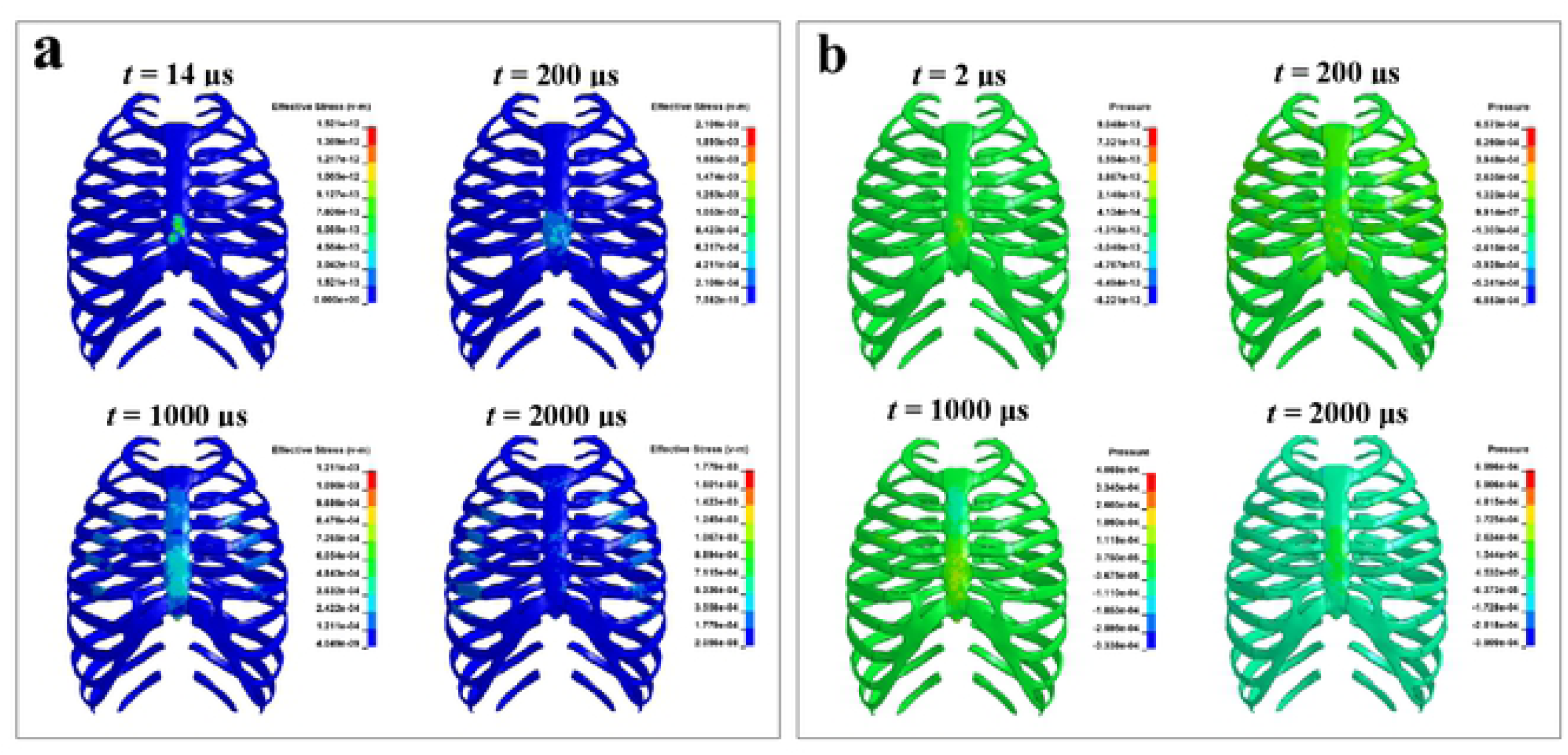
Distributions of (a) stress and (b) pressure of skeleton at different time.

### 3.3. Mechanical response of human organs

The position that the human organs closest to the impact point is selected as the measuring point, and the stress, pressure, acceleration and velocity curves of skeleton and internal organs are shown in Fig. 12 and Fig. 13.

**Fig. 12.**
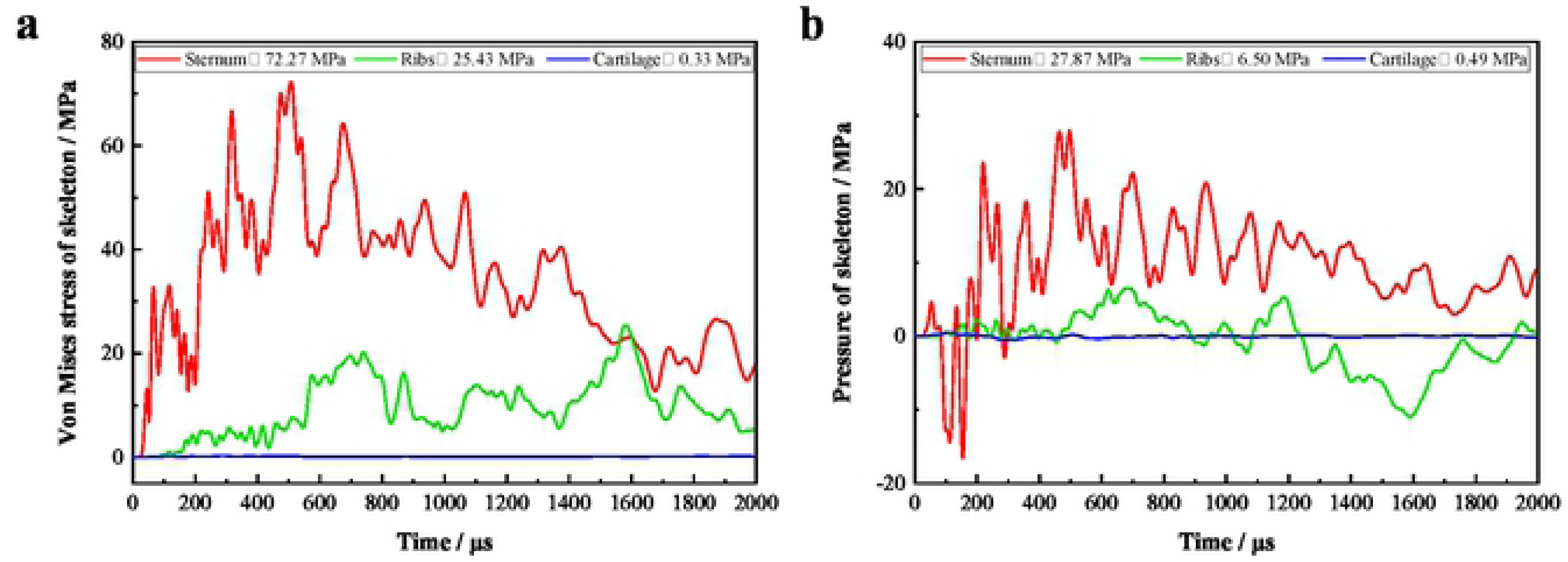
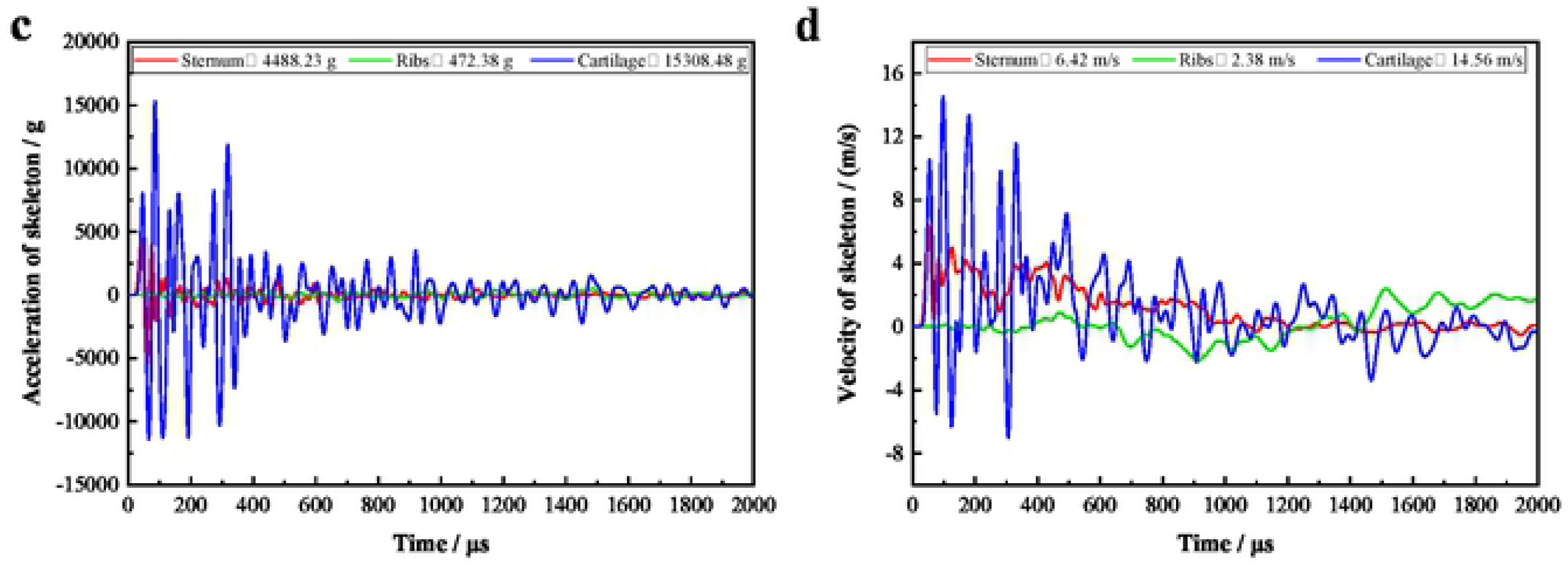
Time histories of (a) stress, (b) pressure, (c) acceleration and (d) velocity of skeleton.

**Fig. 13.**
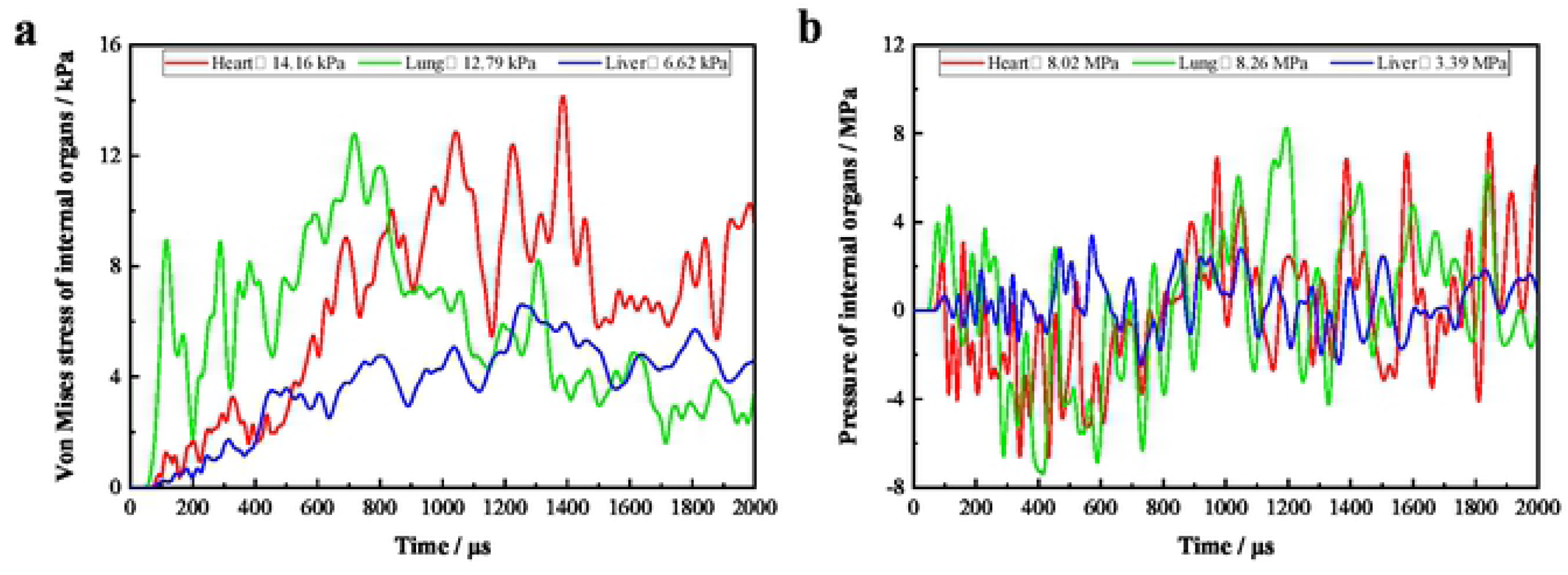
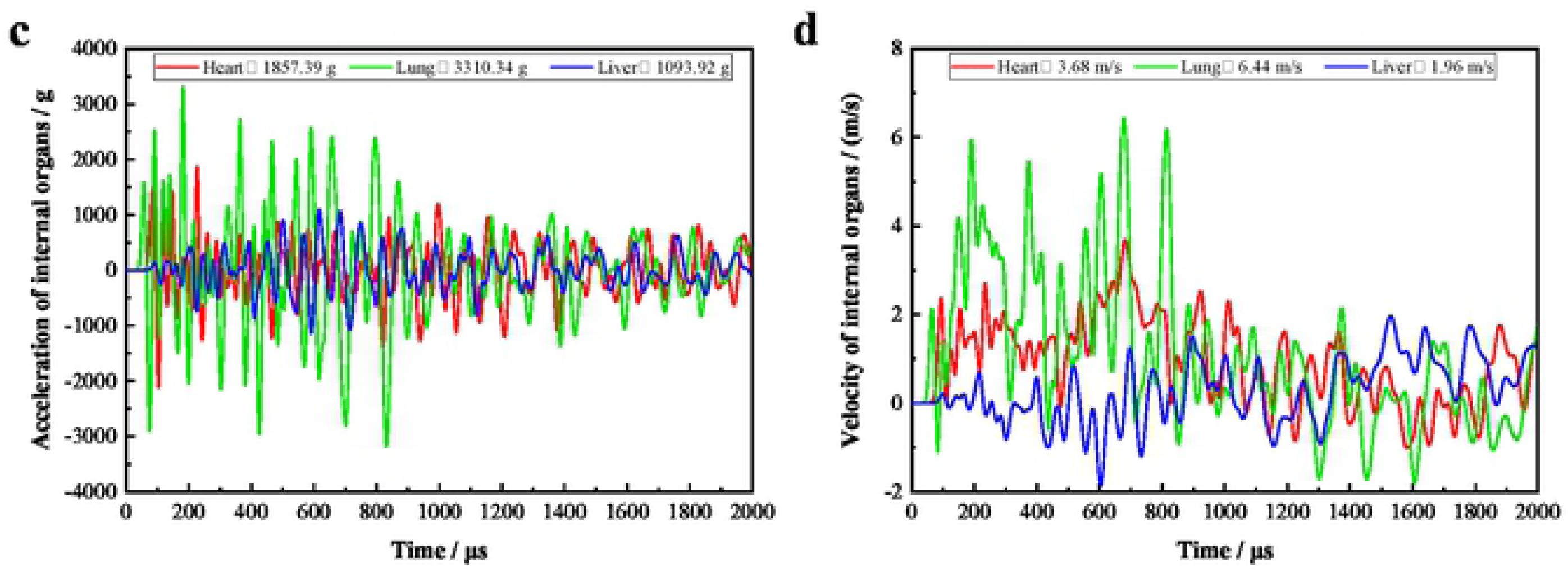
Time histories of (a) stress, (b) pressure, (c) acceleration and (d) velocity of internal organs.

When there is no buffer layer, the mechanical response parameters of various organs are shown in Table 6. The maximum stresses of sternum, ribs, costal cartilage, heart, lung and liver are 72.27 MPa, 25.43 MPa, 0.33 MPa, 14.16 kPa, 12.79 kPa and 6.62 kPa, respectively, and the maximum pressures are 27.87 MPa, 6.50 MPa, 0.49 MPa, 8.02 MPa, 8.26 MPa and 3.39 MPa, respectively. It can be found that the skeleton stress is MPa level, but the internal organs stress is kPa level. In the evolutionary history of human skeleton, the skeletal plays a supporting role in protecting the internal organs such as the lungs and heart, so the skeletal system bears most of the energy. Since the sternum is closest to the impact point, followed by costal cartilage and ribs, the stress and pressure of sternum are greater than that of costal cartilage and ribs. At the same time, the costal cartilage is softer than sternum and ribs, so its pressure and stress are less than sternum and ribs. According to the literature [37], the threshold stress of sternal fracture is 75∼137 MPa, and the simulated peak stress of sternum is 72.27 MPa, so the BCS will not create sternal fracture.

The stress and pressure of lungs are the largest because the lungs are closest to the skeleton and impact point. Relative to the lungs, the heart is located between the two lungs and is far from the impact point. Under the protection of skeletons and lungs, the stress and pressure of heart are small, so the heart injury is also light. If the heart is exposed to excessive impact energy, it will also produce blunt rupture, resulting in serious injury or even death. Moreover, the blood pressure in the heart will be affected by the impact of pistol cartridge and rapidly increase, resulting in bleeding of the heart. The stress and pressure of liver are the smallest because it is farthest from the impact point, and the peak stress is 6.62 kPa. According to the literature [38], the threshold stress of liver rupture is 127∼192 kPa, so the liver does not rupture.

The mechanism of BABT is related to the pressure wave and stress wave generated by the instantaneous deformation of BCS, and on the other hand, it is created by the energy transmitted to the human by BCS [39]. The pressure wave can cause impact injury to the human organs near the impact point. If it is transmitted to the brain through blood, vertebrae and hypodermis, it will bring about indirect brain injury. However, the stress wave can create laceration of human organs. Meanwhile, the acceleration caused by the instantaneous deformation of BCS is also a key factor of human injury. The maximum accelerations of sternum, ribs, costal cartilage, heart, lung and liver are 4488.23 g, 472.38 g, 15308.48 g, 1857.39 g, 3313.34 g, and 1093.92 g, respectively. According to Newton’s second law of motion, when the mass of organ is constant, the greater the peak acceleration, the greater the force on organ, and the greater the degree of human injury. Thus, the costal cartilage in the skeletal system is the most traumatic, and the lung in the internal organs is the most serious [40]. If the threshold acceleration of human injury is exceeded, the internal organs will be displaced, resulting in strain and tear on the contact surface of organs.

Axelsson [41] believed that human injury induced by blast wave was that the chest wall produced a certain inward velocity and compressed the lungs to create severe lung injury. Consequently, the maximum inward chest wall velocity was used as the evaluation index of human injury, and the blast injury model was proposed. The human injury level was divided into five sections: no injury (0.0∼3.6 m/s), trace to slight (3.6∼7.5 m/s), slight to moderate (4.3∼9.8 m/s), moderate to extensive (7.5∼16.9 m/s), >50% lethality (>12.8 m/s). The essence of BABT is precisely because the bulletproof structure will have a certain blunt ballistic impact on human body under the impact of pistol cartridge, resulting the chest wall to move inward and compressing the thoracic organs to cause injury. Hence, the Axelsson injury model can be used as the main evaluation index for BABT. The sternum at the impact point is selected as the velocity of chest wall, and the maximum inward velocity of sternum is 6.42 m/s, so the injury level caused by BCS is slight to moderate. Meanwhile, the difference in the maximum velocities of ribs, costal cartilage, heart, lung and liver are 2.38 m/s, 14.56 m/s, 3.68 m/s, 6.44 m/s, 1.96 m/s, respectively, which also cause injury on the interface of human organs.

The peak pressures at the middle of front of heart and liver obtained by Roberts [18] are 0.743 MPa and 0.130 MPa, respectively, while the peak pressures of heart and liver from the nearest impact point are 8.02 MPa and 3.39 MPa in this paper. It can be seen that the results obtained in this paper are larger than those obtained by Roberts [18]. The main reasons for the differences are: (1) the effect of type and initial velocity of pistol cartridge. Roberts’s research shows that the peak pressure of organs increases with the increase of the velocity of pistol cartridge. (2) The effect of structural features, materials and thickness of body armor. The soft body armor itself can play a certain buffering effect, thus reducing the peak pressure of organs. (3) The effect of the measuring point’s position. Roberts selects the middle of front of organs for measuring point, but the measurement point may not be the point closest to the impact point. For example, the point of liver closest to the impact point is located near the inferior vena cava, and the middle of the front of liver is about 4 cm from the inferior vena cava, resulting in less pressure of organ. (4) The effect of the structural characteristics of human model, such as thicker skin and muscle can play a better buffering role, thereby reducing the injury of internal organs.

### 3.4. Effect of buffer layer on BABT

The main function of buffer layer is to weaken the blunt impact of BCS on human, and reduce the BABT caused by BCS. In practical applications, low-density foam materials such as EPE are often used as buffer layers, and its thickness has a reasonable range [42]. The buffer layer is too thin to protect the human by reducing the effect of BABT. Although increasing the thickness of buffer layer can effectively reduce the BABT, it will increase the weight of BCS and reduce its comfort. Consequently, the buffer layer with thicknesses of 1.0 mm, 2.5 mm and 5.0 mm were selected to study the effect of buffer layer on the mechanical response of human organs. Fig. 14 visually shows the effect of buffer layer thickness on human injury.

**Fig. 14.**
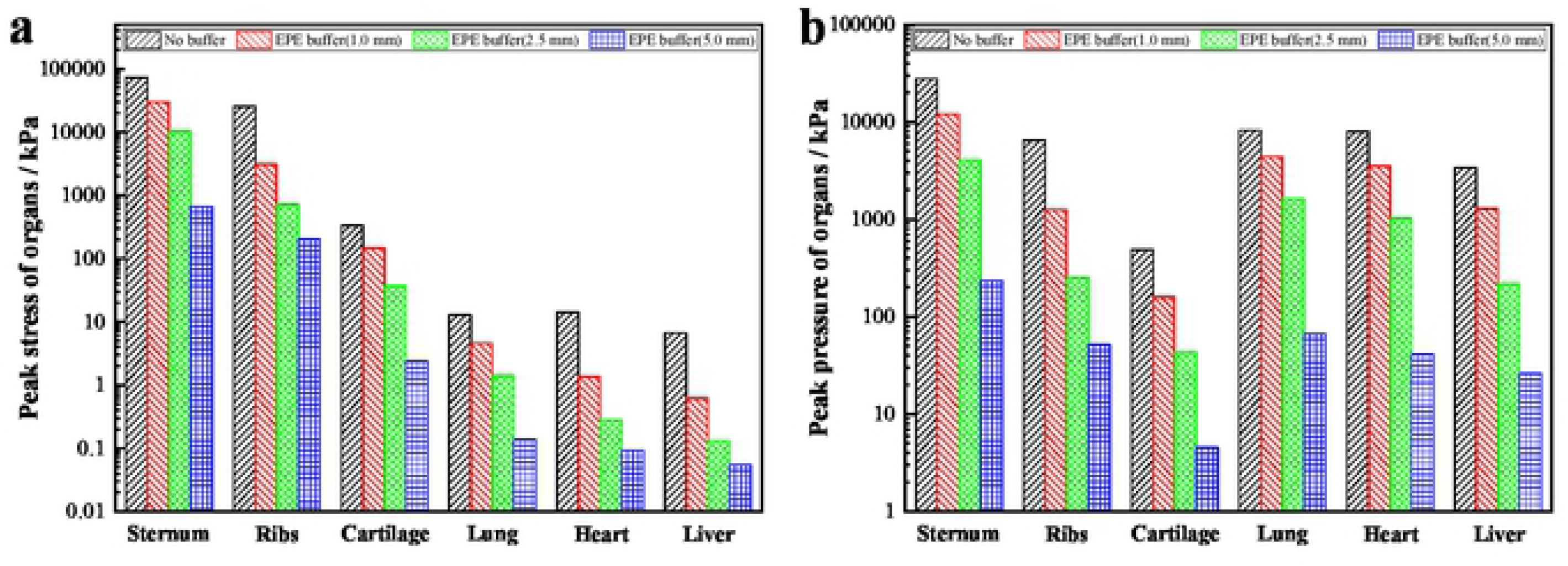
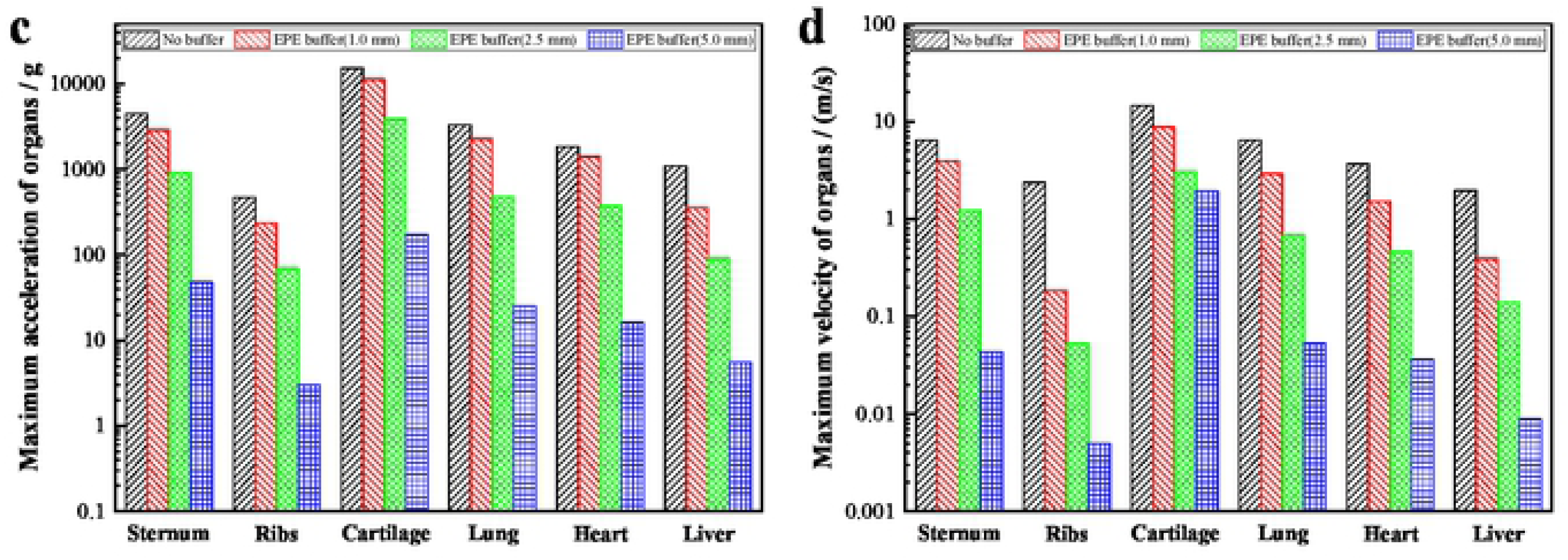
Effects of buffer layer on (a) peak stress, (b) peak pressure, (c) maximum acceleration and (d) maximum velocity of various organs.

The mechanical response parameters of various organs under different thicknesses of buffer layer are shown in Table 7. The sternal stresses corresponding to the EPE buffer layers with thicknesses of 1.0 mm, 2.5 mm and 5.0 mm are 29.95 MPa, 10.16 MPa and 0.66 MPa, respectively, the liver stresses are 0.61 kPa, 0.13 kPa and 0.05 kPa, the sternal velocities are 3.94 m/s, 1.23 m/s, 0.04 m/s. According to the threshold stress of sternum and liver injury, sternal fracture and liver rupture are not caused when the buffer layer is added. According to the Axelsson injury model, the injury level is trace to slight when the thickness of buffer layer is 1.0 mm, while the buffer layer with thickness of 2.5 mm and 5.0 mm will create subtle BABT. By comparing the human injury with or without the buffer layer, it is concluded that the buffer layer can significantly reduce the BABT, and the reduction is related to the thickness of buffer layer, that is, the thicker the buffer layer, the lighter the BABT. Therefore, selection of an appropriate buffer layer is very important for reducing the BABT. Although the buffer layer can reduce the BABT to a certain extent, it is difficult to completely eliminate the BABT [43].

## 4. Conclusion

By simulating the impact process of the pistol cartridge on BCS and the mechanical response process of the human organs, the mechanism of BABT and the effect of the buffer layer on BABT were analyzed. The following conclusions are drawn:

1. The BCS can resist the shooting speed of the 515 m/s without being broken down, and the maximum depression depth of skin at the impact point is 4.98 mm, which is less than 25 mm specified by the Chinese standard and 44 mm specified by the NIJ standard.
2. The maximum energy absorbed by human torso is 37.02 J, which accounts for 5% of the initial energy of pistol cartridge. Although the BCS is not broken down, the energy of pistol cartridge is transmitted to the human torso, thus causing BABT to human organs.
3. The stress waves produced during the impact process begin to propagate from surface to interior, and from near to far around the impact point. Taking the impact point as the center with peak stress value, the stress wave decreases continuously during the process of transmission. Therefore, the closer to the impact point, the more serious the human organ injury.
4. When there is no buffer layer behind the BCS, the peak stresses of sternum and liver are 72.27 MPa and 6.62 kPa, respectively. The BCS will not cause sternal fracture and liver rupture based on the threshold stress of human organs injury. According to Axelsson injury model, the maximum inward chest wall velocity is 6.42 m/s, which creates slight to moderate injury.
5. The injury level is trace to slight when the thickness of the buffer layer is 1.0 mm, while the buffer layer with the thickness of 2.5 mm and 5.0 mm will cause subtle BABT. Therefore, the use of the buffer layer could effectively reduce the BABT. Meanwhile, the thicker the buffer layer, the lighter the human injury. Consequently, it is necessary to provide a buffer layer between body armor and human body for weakening the blunt impact of body armor on human to reduce the BABT.

## Acknowledgements

This work was supported by the National Key R&D Program of China (Grant number 2016YFC0802800 and 2018YFC0809903) and National Natural Science Foundation of China (Grant number 51874041).

